# Multiple goal pursuit – to kill two birds with one stone or to fall between two stools ?

**DOI:** 10.1101/038919

**Authors:** E.Z. Meilikhov, R.M. Farzetdinova

**Affiliations:** NRC “Kurchatov Institute”, 123182 Moscow, Russia; Moscow Institute of Physics and Technology, 141707, Dolgoprudny, Russia

## Abstract

We present the simple phenomenological (but - analytic) model allowing to formalize description of multitasking, i.e. simultaneous performing several tasks. That process requires distribution of attention, and for great number of goals do not lead to success. Our consideration shows that simultaneous performing more than two tasks is, most likely, impossible.

---

It is well known that attention is a limited cognitive resource [1, 2]. That is why performance and attention are related by the simple dependence - decreasing attention span for some task lowers the quality of its performance [3]. That effect is evident in the situation when two, three or more tasks are performed simultaneously. In that case, the performance of each task is derived at the expense of another one [4]. One cannot to multitask without lowering qualities of performing [5].

Distributing attention is the human power to concentrate simultaneously on several objects that provides possibility to perform several actions together. However, there are grounds to believe that at every instant one type of conscious activity could only be accomplished, and the subject feeling of simultaneous performing several tasks stems from fast and frequent shifts between them [6]. With that “simultaneous” performing of two tasks (none of which is automatic one), the performance efficiency of each of them reduces as a result of competition for attention resources [7].

The well-known example of multitasking is a telephoning in the process of driving. Driving performance is significantly degraded by cell phone conversations. In particular, brake reaction times are delayed and accident rates increase [8, 9, 10]. It has been demonstrated that cellphone conversations cause drivers to fail to see up to half of the information in the driving environment that they would have noticed had they not been conversing on the phone [10]. USA National Safety Council estimated that about 30% of all accidents on U.S. highways are caused by drivers using cell phones [2].

In the present work, we try to formalize the description of the multitasking process in the framework of some phenomenological (but analytical) theory. Let us consider the situation, when some fixed proportions *r*′, *r*″ = 1 – *r*′ of processing resources (for instance, attention) are provided for each of two tasks. Degrees *p*′, *p*″ of fulfillment of these tasks depend on the distribution of processing resources: for instance, *p*′ = 1 is the hundred per cent performance of the task I, *p* = 0 is the task I was not performed at all. In the general case, each of tasks is performed in part only (the performance degree varies from 0 up to 1).

Performance-resource functions *p*′ = *p*′(*r*′), *p*″ = *p*″(*r*″) themselves depend on the character of tasks performed. As a rule, they are not known but, in principle, could be experimentally derived. However, it is clear that for “simple” tasks those functions rise monotonously with increasing attention resource and approach asymptotically unity at *r*′, *r*″ → 1. Besides, it is not improbable that there are some resource thresholds 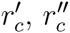 (i.e. some non-zero minimal resources) which are required for corresponding tasks to be performed at least partially. That could be exemplified by the known experiment with “invisible gorilla” [11], which is not visible just owing to the smallness of the resource accessible for its “detection” (*r* < *r_c_*). In general, 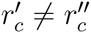 but now we consider only the particular case of simultaneous performing uniform tasks with 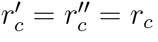.

It is convenient to describe the result of the joint performing of two tasks (I and II) by the diagram like that shown in Fig.1 [3, 12]. Coordinates *p*′, *p*″ of each point on the diagram curve (corresponding to some distribution of processing resources) define probabilities of performing both tasks [13]. As for dependencies *p*″ = *p*″(*p*′) they are itself defined by performance-resource functions. For reasons of mathematical simplicity, it is convenient to approximate that function (for task I) by the simple trigonometric “trial” function

**Figure 1.**
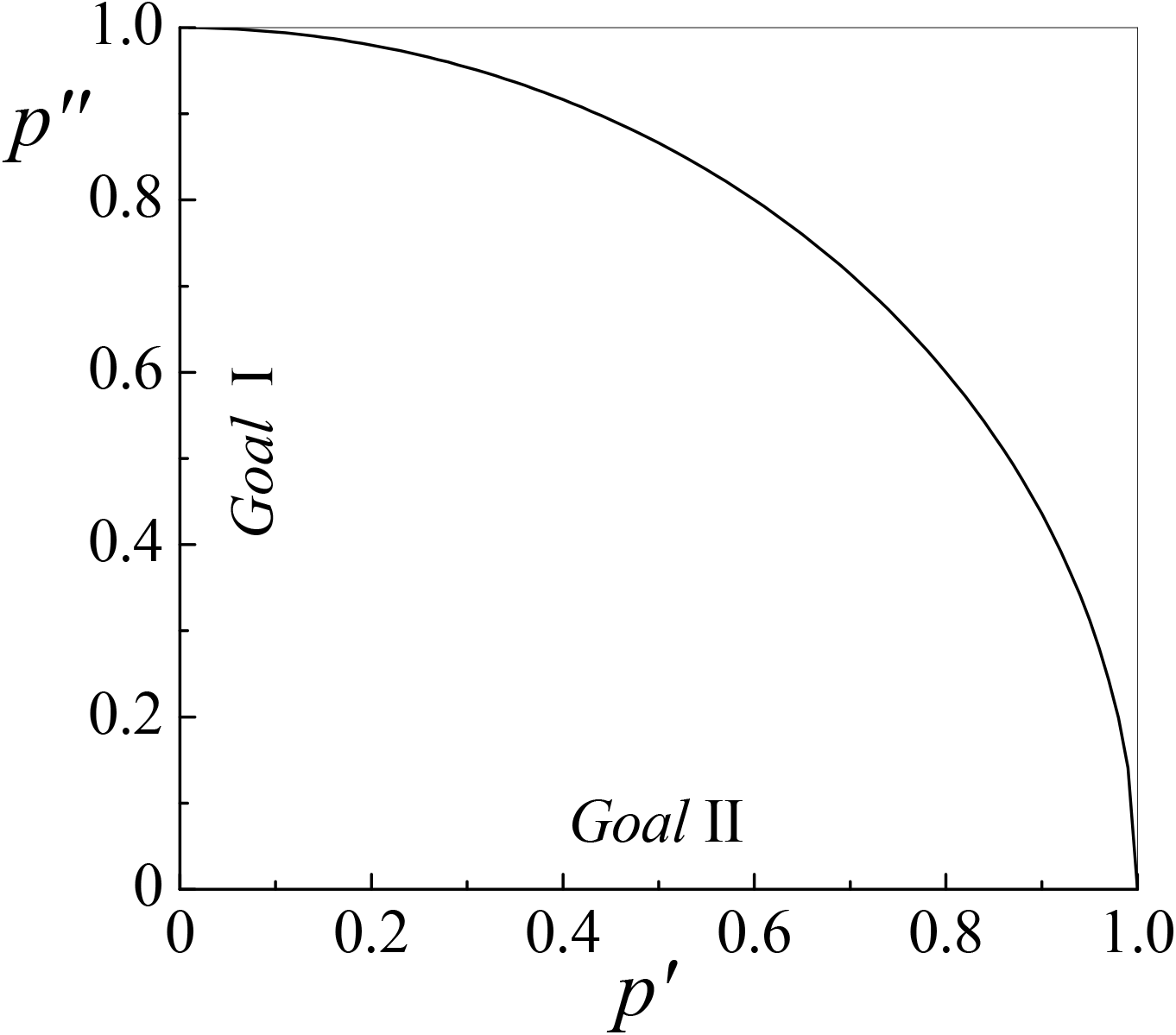
Diagram of simultaneous performing two tasks (I and II) [12].

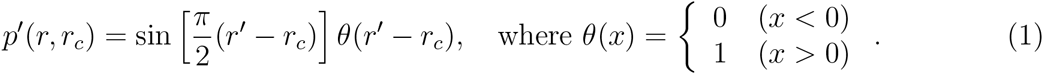

Function *p*′ (*r*,*r_c_*), defined in such a way, is met all above mentioned conditions (its graph is represented in Fig. 2a). Let for simplicity both tasks I and II are characterized by identic performance-resource functions. Then, in the light of *r*′ + *r*″ = 1, we have for task II

**Figure 2.**
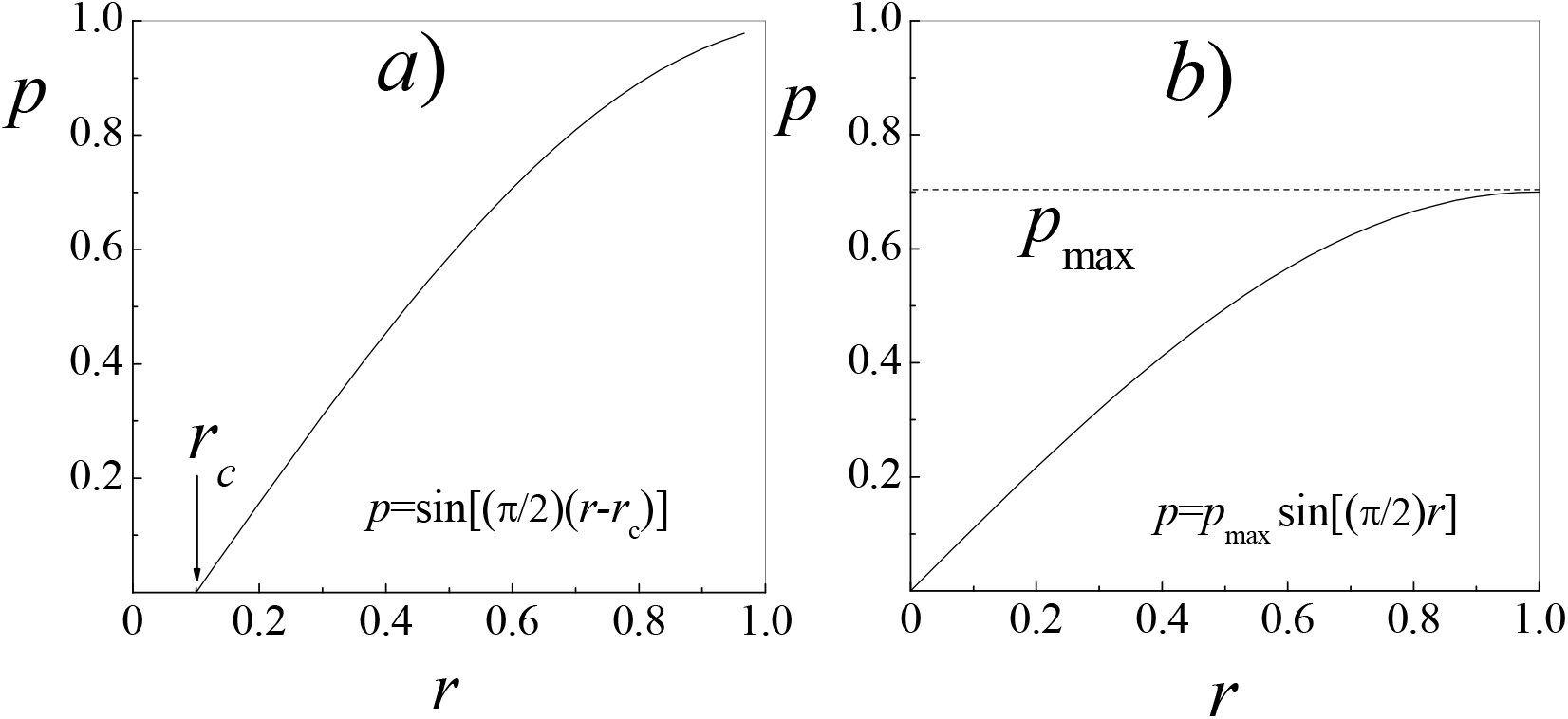
Trial performance-resource function.

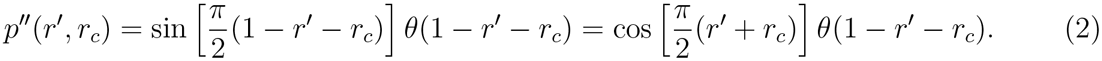

For different thresholds *r_c_*, two relations (1), (2) are parametric representations of corresponding performance-performance dependencies *p*″ = *p*″(*p*′), shown on diagram of Fig. 3. It is seen that at small thresholds there are regimes for which degrees of performing both tasks are rather high (50-70%). Definitely, that is not 100%, but for a variety of tasks even the result of about 20-30% is quite reasonable (for example, amateur fishing, text recognizing during opera audition, etc.).

**Figure 3.**
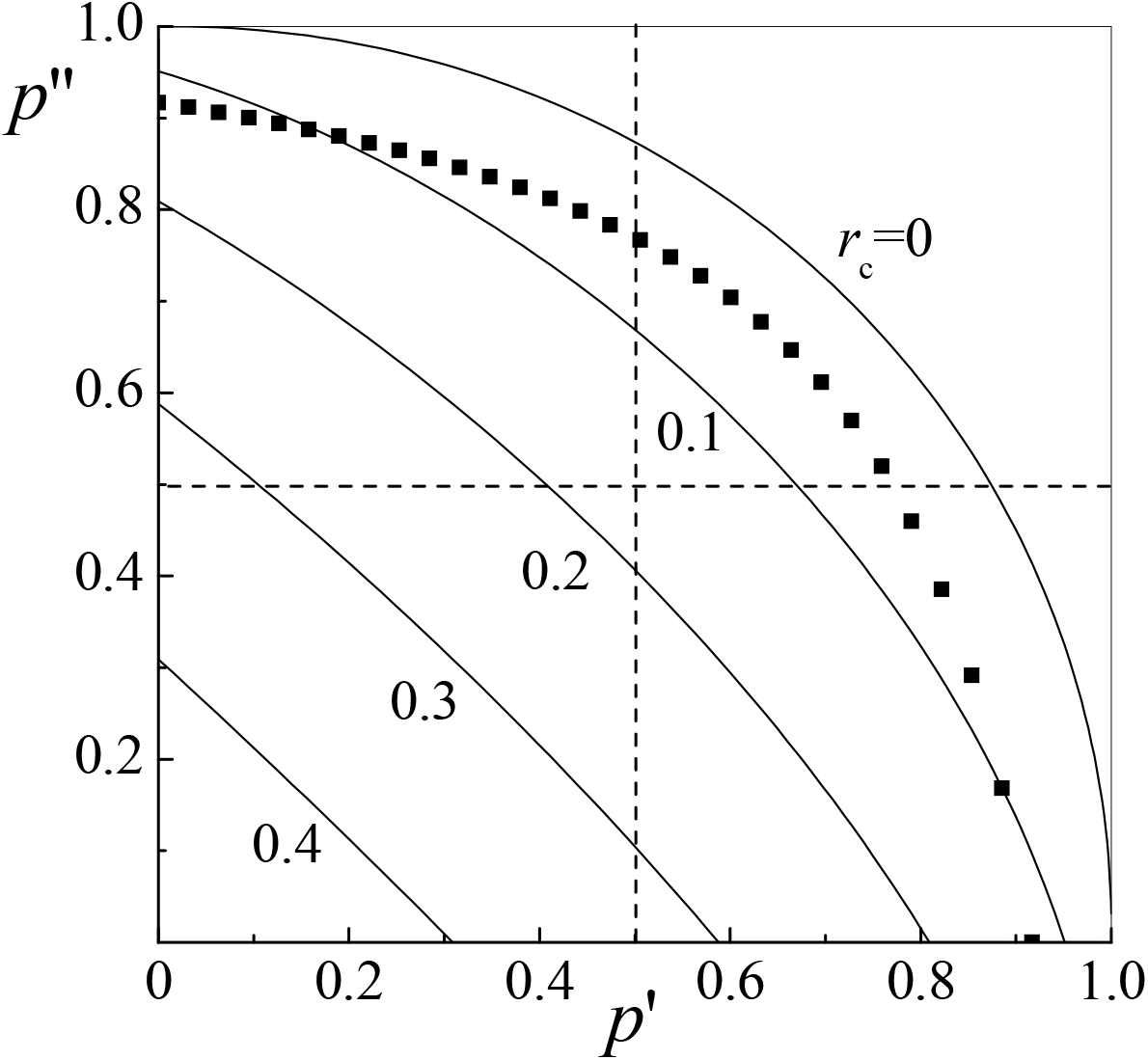
Diagrams of interconnecting parameters *p*′ and *p*″ (performance-performance dependencies) for the performance-resource function (1). Points are the same for the performance-resource function (11).

One can readily see, that for non-threshold tasks (*r_c_* = 0) the dependency *p*″ = *p*″(*p*′) corresponds to the circle (*p*′)^2^ + (*p*″)^2^ = 1. Such a form of the discussed dependency has commonly been suggested (for simplicity) previously without any justification [3, 12]. With non-zero threshold (*r_c_* > 0), the plot of that dependency turns into ellipse. This is clear from the explicit expression *p*″ = *p*″(*p*′) that could be obtained by eliminating *r*′ from Eqs. (1), (2):

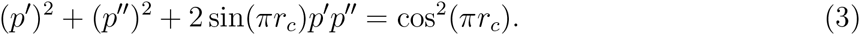

This is the equation for the ellipse with the center in the origin, whose big axis is turned (counter-clock-wise) by an angle of 45° relative to the axis *p*″. From Eq. (3), it follows that coordinates of the point with *p*′ = *p*″ (symmetric, or optimal, distribution of resources) are

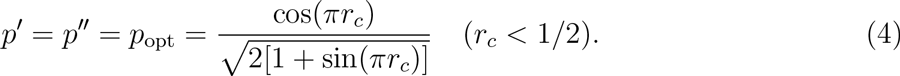

Performance degree, predicted by Eq. (4), declines almost linearly with increasing the threshold *r_c_* (cf. Fig. 4) and amounts to *p*_opt_ ≈ 70% for non-threshold situation (*r_c_* = 0), *p*_opt_ ≈ 38% at *r_c_* = 0.25 and tends to zero at *r_c_* → 0.5.

**Figure 4.**
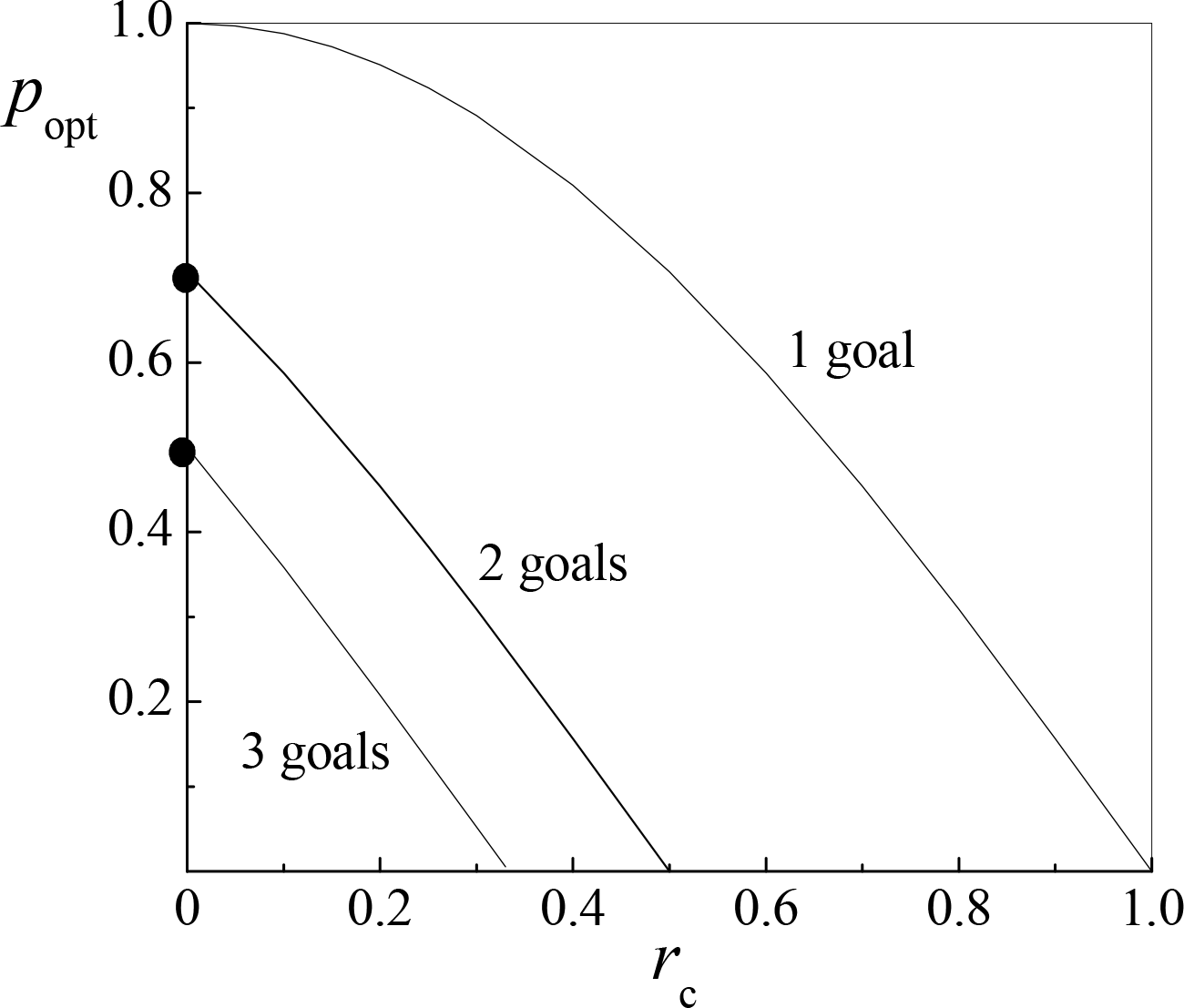
Optimal degree of performing one, two and three tasks for different thresholds *r_c_* of the performance-resource function.

In the case, when thresholds 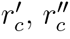 of two tasks are different, the relation (4) remains valid for symmetrical situation (*p*′ = *p*″) with the only substitution 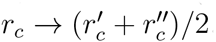.

For concrete tasks, there is always some condition *p* > *p_c_* which has to be met by *both* values *p*′, *p*″ in order for both tasks could be considered as being in the least performed. To take that into account, two dashed lines have been drawn in Fig. 3, corresponding to values *p*′ = *p*″ = *p_c_* with *p_c_* = 0.5. Those lines divide the diagram into four square regions, and those points only correspond to the successive process that enter in the right upper square. Other points correspond to regimes for which or the first task is not performed (the right lower square), either the second one (the left upper square). Surely, we have to account that the criterium of success depends on the concrete type of a task. For instance, one could consider two tasks - (i) to watch a World Cup final and (ii) to stroke a cat - successively performed, if *p*′ ≈ 0.9, *p*″ ≈ 0.1.

In the model framework, it is easy to consider the case, when performance degrees *p*′, *p*″ are limited by the quality of base data (the noise level, for example) or by the limited capacity of the subject himself which does not allow to reach hundred-per-cent performing the task. If maximum attainable levels fall down to *p*′ = *p*″ = *p*_max_ < 1 (cf. Fig. 2b), then values *p*′, *p*″ for symmetrical distribution of resources decrease down to levels *p*′*p*_max_, *p*″*p*_max_. Due to the approximately linear dependency *p*′, *p*″ on the threshold *r_c_*, this means that the curve for *r_c_* = 0.1 in Fig. 3 corresponds to the value *p*_max_ = 0.9, the curve for *r_c_* = 0.2 - to *p*_max_ = 0.8, and so on. Accordingly, the possibility to perform simultaneously and successively two tasks decreases.

Conducted consideration is easily extended to the case of simultaneous performance of more than two tasks. For example, for three tasks we have (bearing in mind that attainable resource for task III equals to *r*′″ = 1 – *r*′ – *r*″)

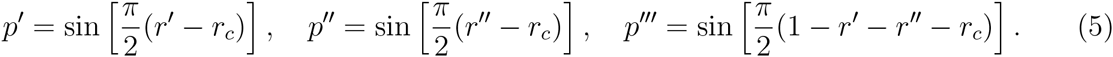

Eliminating parameters *r*′ and *r*″ from those relations, one could find the functional dependency *p*′″(*p*′,*p*″). The latter corresponds to two-dimensional surface whose points have coordinates, defining acceptable triples (combinations) of three parameters *p*′,*p*″,*p*′″. We are restricted by considering the “optimal” variant, when one pays equal attention to every task and performance degrees for all three tasks are equal (*r* = 1/3 and *p*′ = *p*″ = *p*′″ = *p*_opt_). In that case, from Eq. (1) we obtain the relation

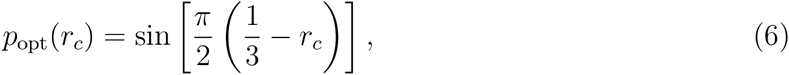

which determines the value *p*_opt_ as a function of the threshold *r_c_*. Specifically, it follows herefrom that in the zero-threshold case (*r_c_* = 0) *p*_opt_ = 1/2, which, as could be expected, is significantly smaller than for simultaneous performance of two tasks only. The whole dependency *p*_opt_(*r_c_*) is shown in Fig. 4.

If one consider the performance of all three tasks on the level of 50% as a success, that, as could be seen, is possible in the zero-threshold regime only. May be, Gaius Julius Caesar did have this property? Indeed, according to legend he could pursue multiple goals concurrently.

Now, actually, we could formulate the theorem:

> *Simultaneous performing more than two tasks is, most likely, impossible.*

In contrast to absolutely faithful mathematical statements, that “theorem” is of some obscure and probabilistic character, that is quite natural for such a non formalized science as cognitive psychology.

The result obtained is directly generalized to the case of arbitrary number *N* of simultaneously performed tasks (when *r* = 1/*N*):

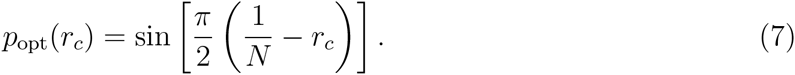

Particularly simple result is obtained, if *r_c_* = 0. In that case, from Eq. (7) it follows

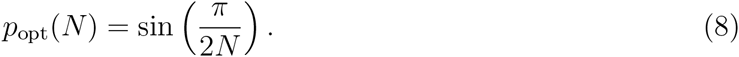

Correspondent dependency *p*_opt_(*N*) is presented in Fig. 5. As could be expected, increasing the number of tasks lowers significantly the degree of their performance.

**Figure 5.**
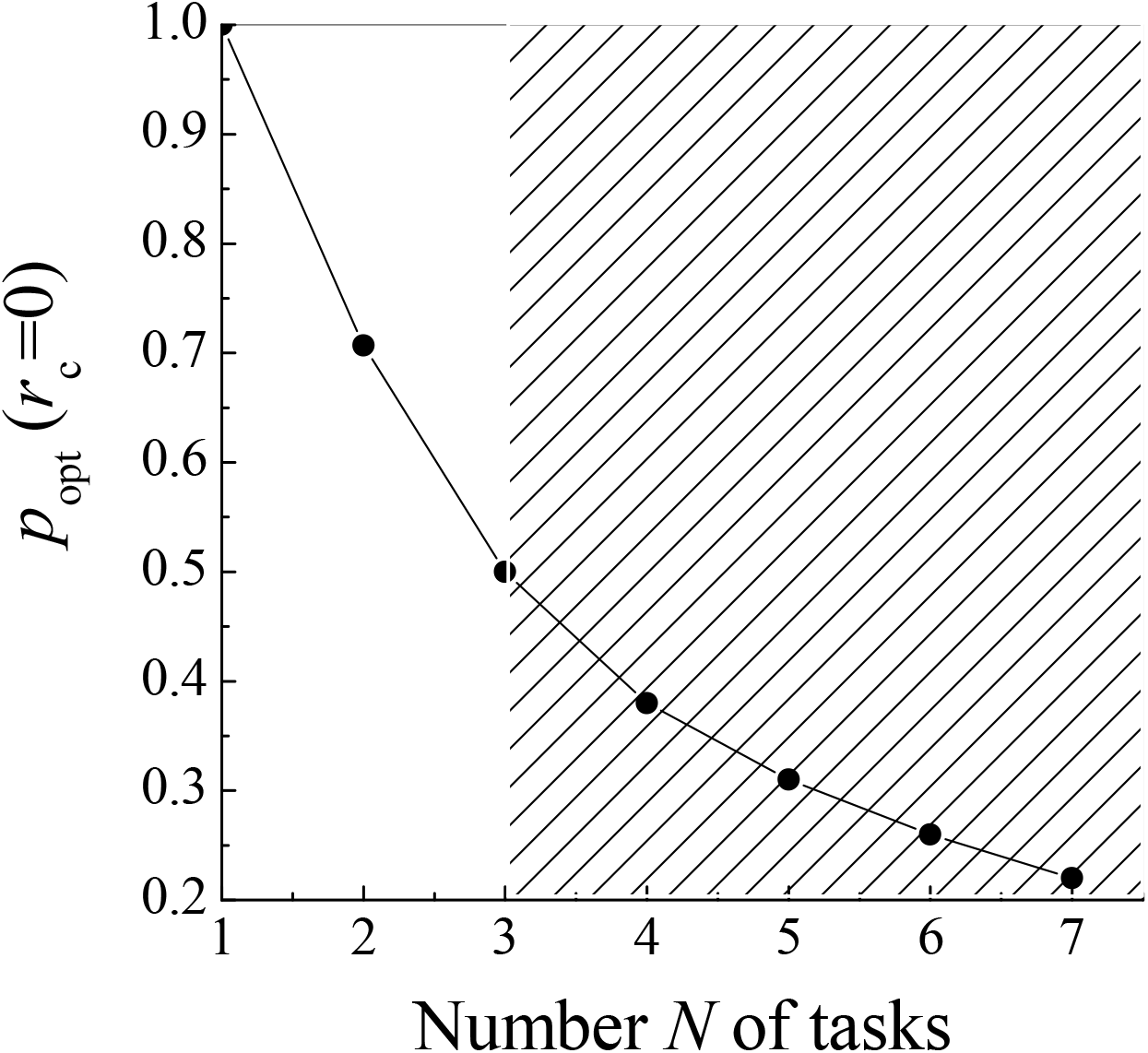
Optimal degree of simultaneous performing *N* zero-threshold tasks.

Various points of *N*-dimensional surface *p*^(*n*)^ = *p*^(*n*)^[*p*′,*p*″…, *p*^(*N*)^] correspond to different variants of simultaneous performing *N* tasks. For the integral estimate of those variants one could introduce the success ratio, determined by the parameter

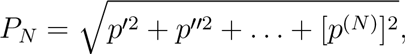

which is the distance between the coordinate origin and the corresponding point *p*′, *p*″,…, *p*^(*N*)^ on that surface. In the “optimal” regime (when *p*′ = *p*″ =… = *p*^(*N*)^ = *p*_opt_ and *r_c_* = 0),

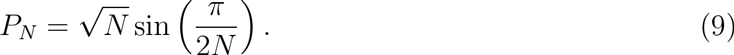

Only for *N* =1 and *N* = 2 we have *P_N_* = 1 (*P*_1_ = 1 - the task is completely performed, *P*_2_ = 1 - both tasks are performed by *p*_opt_ ≈ 70%). For 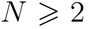, always *P_N_* < 1 (for instance, 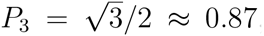, *P*_4_ = 2sin(*π*/8) ≈ 0.76 and so on). That result confirms the above-formulated theorem once more.

Similar results could be also obtained for other reasonable performance-resource “trial” functions *p*′ = *p*′ (*r*′):

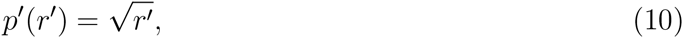

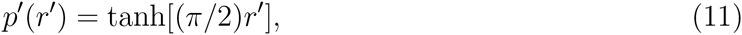

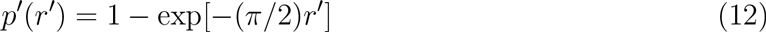

(cf. [13]). For the function (10), the performance-performance dependency for two tasks is defined by the same “circle” relation *p*′^2^ + *p*″^2^ = 1 as for zero-threshold (*r_c_* = 0) trial function (1).

At the same time, for the function of the type (11) the performance-performance dependency for two tasks is defined by the relation

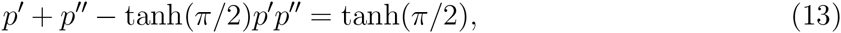

to which corresponds not the circle but the hyperbola with the symmetry axis making the angle 45° with the axis *p*′. Nevertheless, as could be seen from Fig. 3, the plot of that hyperbola (represented by points) is very close to the zero-threshold (*r_c_* = 0) circle dependency *p*″ (*p*′).

That result is the evident consequence of the plot similarity of functions (1) and (11) in spite of their different analytic representations.

Those functions correspond to the “simple” tasks, for which *p*′ (1) ≈ 1. The last condition means that if one pays the full (or almost full) attention to a single task, then it is practically performed in each case. For “complex” tasks the situation is not the same, and the form of the performance-resource function itself, which is defined experimentally [3], could provide the quantity estimate for the concept of the “task complexity”. Let, for example, the performance-resource function is approximated by the dependency

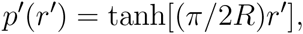

where the parameter *R* could be of any value (the trial function (11) corresponds to *R* = 1). Then the task performance will be highly successive (*p*′ (1) ≈ 1) at *R* << 1 and, conversely, less successive (*p*′(1) << 1) at *R* >> 1. Hence, the parameter *R* value is the measure of the task complexity: *R* << 1 - the task is “simple”, *R* >> 1 - the task is “complex”, *R* ≈ 1 - the task is of the “intermediate” complexity.

Actually, the task complexity is rather relative concept: what is complex for one man, might be simple for another. Moreover, the task complexity level could change in the course of learning and training [14]. Thereby occurring changes of the performance-resource function of Eq. (11)-type are shown in Fig. 6: with training, maximum of that function becomes to be closer to the value *p* ≈ 1, corresponding to the full task performing. Formally, that is the consequence of gradual reducing the complexity parameter *R.*

**Figure 6.**
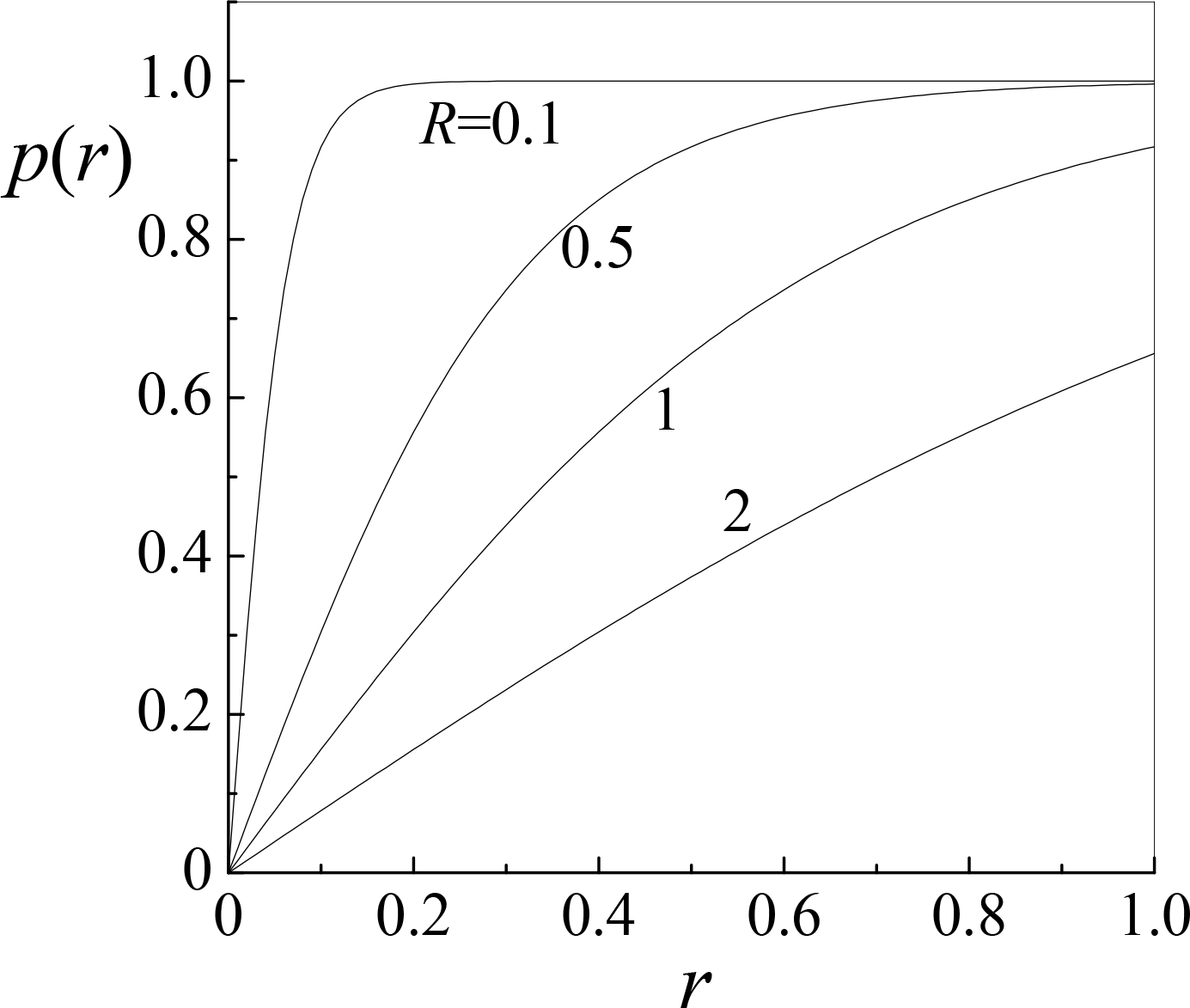
Performance-resource functions (11) for tasks of different complexity R.

For the simultaneous performing two tasks of different complexities *R*_1_ and *R*_2_ performance-resource functions have the form

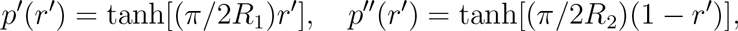

wherefrom it follows for performance-performance function

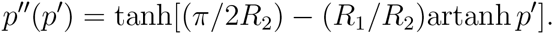

Plots of those functions for the case, when one of the tasks is of the intermediate complexity (*R*_1_ = 1) and another one is more complex (*R*_2_ >> 1) or more simple (*R*_2_ << 1) than the former, are represented in Fig. 7. Naturally, possible degree of performing the second task increases with its simplifying. As a quantitative estimate for the success of the simultaneous performing two tasks one could consider the maximum value 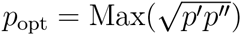 of the geometric mean of tasks’ performing degrees *p*′, *p*″. With fixed complexity of the first task (*R*_1_ = 1) that maximum value depends on the complexity of the second task. Corresponding optimal regimes are marked by points on graphs of Fig. 7. We see that *p*_opt_ ≈ 0.2 at *R*_2_ = 10, *p*_opt_ ≈ 0.7 at *R*_2_ = 1 and *p*_opt_ ≈ 0.9 at *R*_2_ = 0.25. The first case does not suitable, but two later cases could be considered as rather satisfactory as far as the simultaneous performing of two tasks is concerned.

**Figure 7.**
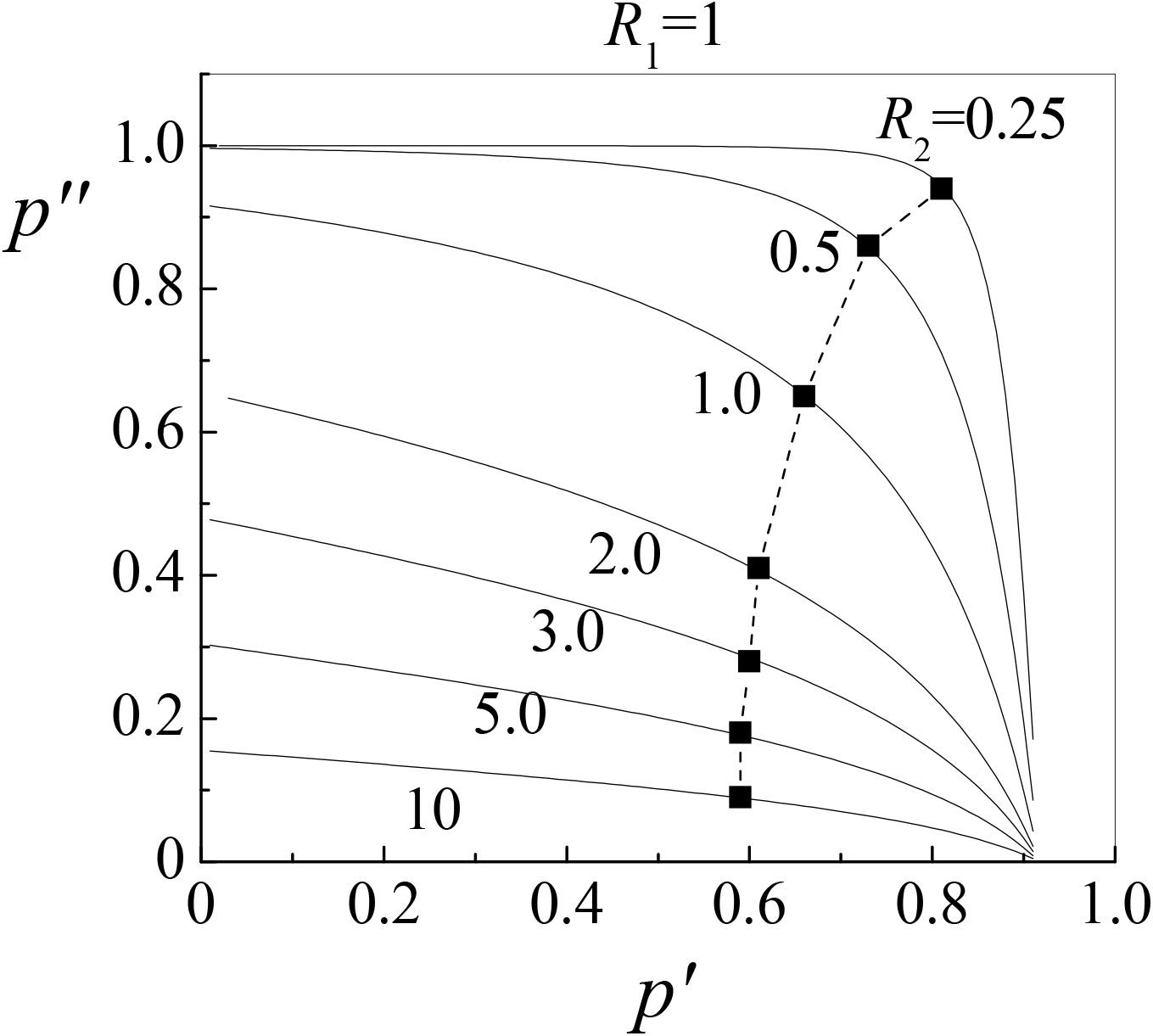
Performance-performance functions for simultaneous performing two tasks of different complexities *R*_1_ and *R*_2_. Optimal regimes are marked by points.

Notice, that, in a sense, the performance-resource function (10) is the simplest one. Independent of the number *N* of simultaneously performed tasks, it determines the performance-performance function, which corresponds to *N*-dimensional hyper-sphere

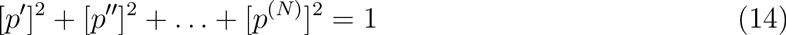

of the unit radius.

That simplicity is kept for some more general threshold function 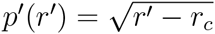 which is also associated with the hyper-sphere

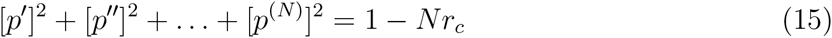

of the radius 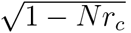, reducing with increasing *N*. Evidently, that corresponds to lowering the performance degree with increasing the task number.

Thus, in the present work we suggest the simple phenomenological (but analytical) model, allowing to formalize considering the process of multitasking, i.e. simultaneous performing several tasks. That process requires distributing attention and rarely results in the success if the number of tasks is large. Our consideration shows that simultaneous performing more than two tasks is, most likely, impossible.

The considered question is only the specific example of the problem associated with the fact that cognitive psychology is practically descriptive science and is not formalized in the mathematical sense. However, according to I. Kant - “In each natural science so much truth is concluded, how many mathematics in it is”. The goal of the work is to show that even such a “non-mathematized” science as the cognitive psychology could be moved to the rank of “true” natural sciences.

We have given the analytical description of the known cognitive multitasking process in the framework of relatively simple mathematical idea. We have aimed not to simulating that psychic phenomenon but to presenting it in the mathematical (analytic) form. For that we have attempted to select the “core” of the process and to cut unimportant details. Therein lies the sense of centuries-long approach to the truth in such “true” sciences as physics or chemistry, where the first place has always belonged to constructing and subsequent extending various (but always - approximate) models. Of course, such an approach never leads to the absolutely accurate description, but allows to grasp the essential features of described phenomena and to construct models whose accuracy is sufficient enough to use them further for solving various practical tasks.

Of course, the problem is not exhausted by the modeling of multitasking. Analytical models of wide variety of cognitive processes could be effective for the improving reliability and operating speed of interfaces brain-computer, ergonomic provision of operator performance within different industrial areas, increasing the level of the creative activity at solving different professional problems.

